# Expression of the ACE2 virus entry protein in the nervus terminalis reveals the potential for an alternative route to brain infection in COVID-19

**DOI:** 10.1101/2021.04.11.439398

**Authors:** Katarzyna Bilinska, Christopher S. von Bartheld, Rafal Butowt

## Abstract

Previous studies suggested that the SARS-CoV-2 virus may gain access to the brain by using a route along the olfactory nerve. However, there is a general consensus that the obligatory virus entry receptor, angiotensin converting enzyme 2 (ACE2), is not expressed in olfactory receptor neurons, and the timing of arrival of the virus in brain targets is inconsistent with a neuronal transfer along olfactory projections. We determined whether nervus terminalis neurons and their peripheral and central projections should be considered as a potential alternative route from the nose to the brain. Nervus terminalis neurons in postnatal mice were double-labeled with antibodies against ACE2 and two nervus terminalis markers, gonadotropin-releasing hormone (GnRH) and choline acetyltransferase (CHAT). We show that a small fraction of CHAT-labeled nervus terminalis neurons, and the large majority of GnRH-labeled nervus terminalis neurons with cell bodies in the region between the olfactory epithelium and the olfactory bulb express ACE2 and cathepsins B and L. Nervus terminalis neurons therefore may provide a direct route for the virus from the nasal epithelium, possibly via innervation of Bowman’s glands, to brain targets, including the telencephalon and diencephalon. This possibility needs to be examined in suitable animal models and in human tissues.

## INTRODUCTION

Many previous reports have suggested that the severe acute respiratory syndrome coronavirus 2 (SARS-CoV-2) gains access to the brain by using an olfactory route from the nose to the brain (Bougakov et al., 2020; Briguglio et al., 2020; Butowt and Bilinska, 2020; Li et al., 2020; Natoli et al., 2020; Meinhardt et al., 2021; Zubair et al., 2021; Burks et al., 2021), similar to some other neuro-invasive viruses that are known to infect olfactory receptor neurons and spread from these first-order olfactory neurons to secondary and tertiary olfactory targets in the brain (Barnett and Perlman, 1993; van Riel et al., 2015; Dubé et al., 2018). Indeed, it has been shown that SARS-CoV-2 can accumulate in various brain regions, in animal models (reviewed in: Butowt and von Bartheld, 2020; Rathnasinghe et al., 2020; Butowt et al., 2021) and in a small number of human patients with COVID-19 (Ellul et al., 2020; Matschke et al., 2020; Meinhardt et al., 2021; Mukerji and Solomon, 2021; Solomon, 2021; Thakur et al., 2021).

However, the route along the olfactory nerve from the nose to the brain is controversial for SARS-CoV-2, primarily for two reasons: (1) the olfactory receptor neurons do not express the obligatory virus entry receptor, angiotensin-converting enzyme 2 (ACE2), or expression is restricted to a very small subset of these neurons (Butowt and von Bartheld, 2020; Cooper et al., 2020; Brechbühl et al., 2021; Butowt et al., 2021). Because sustentacular cells tightly enwrap olfactory receptor neurons (Liang, 2020), these ACE2-expressing support cells can easily be mistaken for olfactory receptor neurons, resulting in false positive identification. (2) The timeline of appearance of SARS-CoV-2 in the brain is inconsistent with a “neuron-hopping” mode: infection of third-order olfactory targets should occur with a significant delay after infection of the olfactory epithelium, as has been reported for other neuro-invasive viruses (Barnett et al., 1995), but instead the hypothalamus and brainstem are reported to be infected as early as, or even earlier than, the olfactory bulb (de Melo et al., 2021; Zheng et al., 2020), and SARS-CoV-2 may even skip the olfactory nerve and olfactory bulb on its way to brain infection (Winkler et al., 2020; Zhou et al., 2020; Carossino et al., 2021). These findings have raised doubt about the notion that the olfactory nerve serves as a major conduit for brain infection in COVID-19 (Butowt et al., 2021).

With few exceptions (Briguglio et al., 2020; Butowt and von Bartheld, 2020; Butowt et al., 2021), studies suggesting an olfactory route for SARS-CoV-2 to achieve brain infection fail to consider the potential for an alternative route from the nose to the brain, the route via the nervus terminalis. Many peripheral processes of the nervus terminalis innervate the olfactory epithelium, the blood vessels below this epithelium, as well as cells in Bowman’s glands (Larsell, 1950), and the central processes of some of these neurons extend to various targets in the forebrain as far caudal as the hypothalamus (Pearson, 1941; Larsell, 1950; Schwanzel-Fukuda et al., 1987; Demski, 1993; von Bartheld, 2004). Some of the nervus terminalis neurons are in direct contact with spaces containing cerebrospinal fluid (CSF) in the region of the olfactory nerve and bulb (Jennes, 1987). About 30-40% of the neurons of the nervus terminalis express gonadotropin-releasing hormone (GnRH), and some of these neurons may release GnRH into blood vessels below the olfactory epithelium (Jennes, 1987; Schwanzel-Fukuda et al., 1987), while other neuronal populations of the nervus terminalis system are thought to regulate blood flow and blood pressure in the nose and forebrain (Larsell, 1918; Oelschläger et al., 1987; Ridgway et al., 1987). These properties make the nervus terminalis a strong candidate for expression of ACE2, which is known to regulate blood flow and blood pressure in many tissues (Tikellis and Thomas, 2012). Expression of ACE2 in the nervus terminalis would suggest that this cranial nerve is a plausible alternative to the olfactory nerve for the SARS-CoV-2 virus to gain access to the brain. However, it has not been previously examined and reported whether nervus terminalis neurons express the obligatory viral entry receptor, ACE2, and any other virus entry proteases such as TMPRSS2 and cathepsins B and L. We have therefore examined whether these entry proteins are expressed in nervus terminalis neurons in an animal model, the postnatal mouse.

## MATERIALS AND METHODS

### Animals and tissue processing

A total of eight wildtype C57BL/6J mice (Jackson Laboratory) at age 3-4 weeks old were used to obtain tissues for experiments. Mice were housed with a 12/12 hours light/dark cycle and given access to water and food ad libitum. All animal experiments were approved by the local ethics committee for animal research at Bydgoszcz (Poland). Immediately after cervical dislocation, the mice were exsanguinated and tissues were dissected. Olfactory epithelium and brain were frozen at −80°C for storage and further usage, or fixed for 3 hours at 4°C in 4% (w/v) freshly prepared paraformaldehyde in phosphate-buffered saline (PBS, pH 7,5), and then incubated in 25% (w/v) sucrose/PBS at 4°C for 16–24 hours, frozen in Tissue-Tek O.C.T. (Sakura Finetek), and cryosectioned at 10-12 μm using a Leica CM1850 cryostat.

### ACE2 −/− knockout (ACE2 KO) control

To verify the specificity of the ACE2 antibody, an ACE2 knock-out (KO) mouse line was obtained from Taconic (strain #18180). Two male homozygous ACE2 KO mice at age 3 weeks old were processed and immunolabeled as described below for wildtype mice. Genotyping was performed according to the manufacturer’s suggested PCR protocol. Lack of an ACE2 protein band was confirmed by using Western blots as described previously (Bilinska et al., 2020). In brief, tissue was homogenized on ice in N-Per Total Protein Extraction reagent (Thermo Scientific) with addition of protease and phosphatase inhibitor cocktails (Sigma-Aldrich). Homogenates were centrifuged for 30 minutes at 20,000 g at 4°C and supernatants were collected. Protein content was measured by the BCA method (Thermo Scientific). Equal amounts of total proteins were mixed with 4x Laemmli sample buffer and boiled for 10 minutes at 80°C. Protein extracts were separated on SDS-PAGE 7.5% gels and mini-protean III apparatus. GAPDH was used as positive control and to verify equal loading. Proteins were blotted to nitrocellulose membranes using standard Tris-glycine wet method. Membranes were blocked with 5% dry milk (Bio-Rad), incubated with goat polyclonal anti-ACE2 (R&D Systems AF3437) at 1/1000 or rabbit polyclonal anti-GAPDH (Protein-Tech, #10494-1-AP) at 1/5000 dilution overnight at 4°C, washed several times in TBST buffer (pH 8.0) and incubated 60 minutes with secondary antibody, anti-goat-HRP (Protein-Tech). Signal was detected using Clarity Max chemiluminescence substrate (Bio-Rad). For confirmation, blots were stripped and re-probed with an additional rabbit monoclonal anti-ACE2 antibody (Abclonal, #A4612). Blots were prepared in three separate experiments with comparable results.

### ACE2 immunocytochemistry and co-localization analysis

For double immunofluorescence labeling, antigen retrieval procedure was performed on frozen sections cut at 10-12 μm. Sections were incubated overnight with a mixture of primary goat anti-ACE2 at 1/500 dilution (R&D Systems, #AF3437) and rabbit anti-GnRH (gonadotropin releasing hormone) at 4°C. On the next day, sections were washed five times in PBST (PBS with 0.05% Triton X-100) and incubated with a mixture of secondary anti-rabbit-AF488 antibody and anti-goat-AF594 at 1/500 dilution for 60 minutes at room temperature. Next, sections were stained for 5 minutes at room temperature in Hoechst 33258 (Sigma-Aldrich) to visualize cell nuclei, and sections were then embedded in aqueous antifade medium (Vector laboratories). Alternating cryosections were incubated with rabbit polyclonal anti-CHAT (choline acetyltransferase) instead of rabbit anti-GnRH antibody in the double staining primary antibody mixture. Occasionally, sections were incubated with anti-OMP (olfactory marker protein) at 1/500 dilution in PBST, following the same protocol. After immunocytochemical reactions, sections were analyzed on a Nikon Eclipse 80i microscope and images were taken using a Nikon DP80 camera. Microscopic images were processed using cellSens Dimension 1.13 software (Olympus). Antibodies and vendors are listed in Supplemental Material, Table S1. To compare the signal intensity between nervus terminalis neurons and cells known to express ACE2 and to internalize SARS-CoV-2, the ACE2 fluorescent signal was compared by measuring the optical density of the signal in gray scale (8-bit maps) in ACE2-expressing nervus terminalis neurons and in sustentacular cells of the dorsal olfactory epithelium, using cellSens Dimension 1.13 software (Olympus). Intensity values were defined by regions of interest and a quantitative immunofluorescence score was calculated by comparing the target mean gray intensity for 15 sustentacular cells and 12 GnRH-positive nervus terminalis neurons.

### Cell counting and statistical analysis

For counting double-labeled neurons, five male wildtype mice at age 3-4 weeks old were used. Approximately every third coronal cryosection (10-12 μm thickness) was stained as described above, and positive neurons were counted in tissue sections under a fluorescent microscope as indicated in Fig. 1 (the medial region from the posterior olfactory epithelium to the caudal end of the olfactory bulb). For each animal, the percentage of double labeled GnRH+/ACE2+ neurons was calculated in relation to the total number of GnRH-positive neurons detected. The same protocol was applied for counting cholinergic nervus terminalis neurons co-labeled with ACE2. A total number of 119 GnRH-positive neurons and a total of 52 CHAT-positive neurons were counted from 3-5 animals, as shown in detail in the Supplemental Material, Table S2. The results were analyzed using GraphPad Prism software. Results are presented as mean ± standard error of the mean (SEM). An unpaired t-test was applied to determine whether the difference in ACE2 colocalization between GnRH+ and CHAT+ neurons was statistically significant. For details of quantification, see the Supplementary Material, Table S2.

**Fig. 1.**
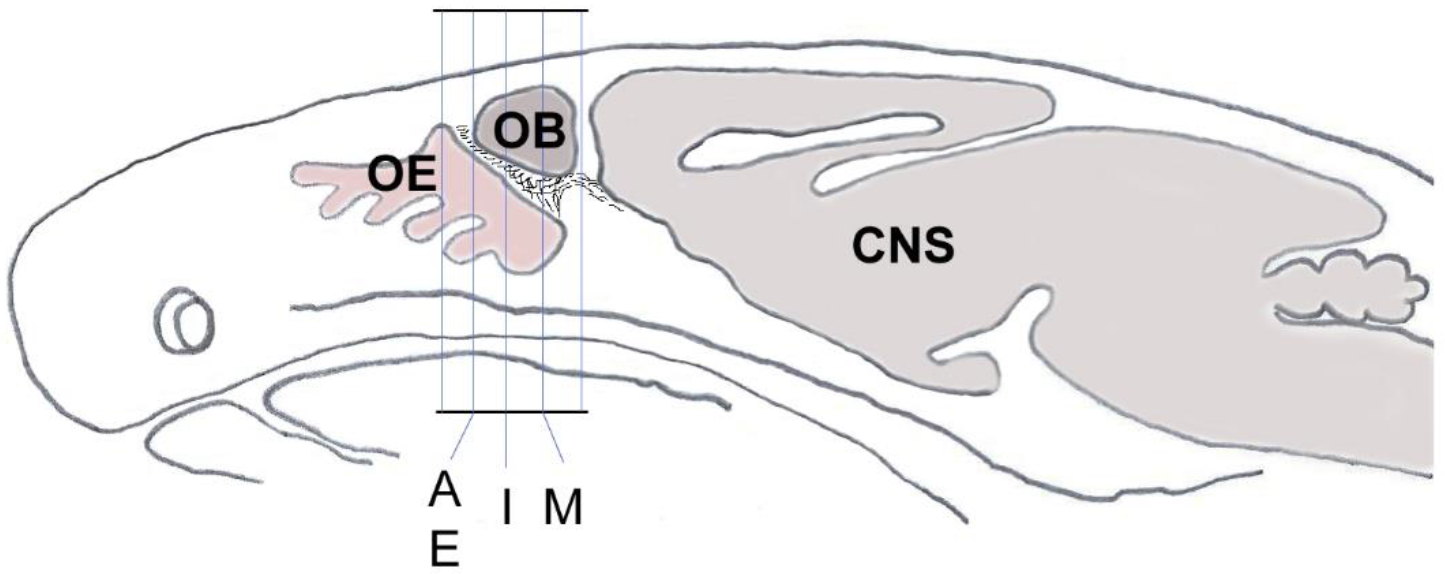
Schematic sagittal section through a mouse head shows the orientation and planes of tissue sections from Fig. 2A, E, I and M. Sections within those planes were used for demonstration of double-immunolabeling and for cell counting. CNS, central nervous system; OB, olfactory bulb; OE, olfactory epithelium.

### TMPRSS2 and cathepsin B and L immunocytochemistry

TMPRSS2 and cathepsins B and L are proteases that SARS-CoV-2 can use to gain entry into host cells (Shang et al., 2020). To determine whether nervus terminalis neurons also express TMPRSS2, tissue sections were incubated with TMPRSS2 antibodies at 1:50 or 1:200 dilution as recommended by the manufacturer. We tested one cathepsin B and one cathepsin L antibody and three different anti-TMPRSS2 antibodies (listed in the Supplementary Table S1) to determine the presence of these proteases in GnRH-positive nervus terminalis neurons. The same protocol as described above for the ACE2 antibody was used. Quantification of GnRH and cathepsin L co-labeled neurons was performed using the same protocol as described for quantification of neurons labeled with ACE2.

## RESULTS

It was previously shown that GnRH is a marker for a major fraction of nervus terminalis neurons (Jennes, 1987; Schwanzel-Fukuda et al., 1987; Demski, 1993; Kim et al., 1999; von Bartheld, 2004). Immunolabeling for GnRH in 3-4 week-old mice showed labeled neurons localized along the olfactory nerve between the olfactory epithelium and the olfactory bulbs (Fig. 2A-H), as expected from previous studies in rodents (Schwanzel-Fukuda et al., 1986; Wirsig and Leonard, 1986; Schwanzel-Fukuda et al., 1987). The majority of the GnRH-positive nervus terminalis neurons was located along the midline in the posterior part of the olfactory epithelium and adjacent to the olfactory bulbs. Preliminary examination revealed that these cells were in the same vicinity as cells labeled with the ACE2 antibody (Fig. 2A-B, E-F). The large majority of GnRH-positive nervus terminalis neurons were fusiform and unipolar in shape.

**Fig. 2.**
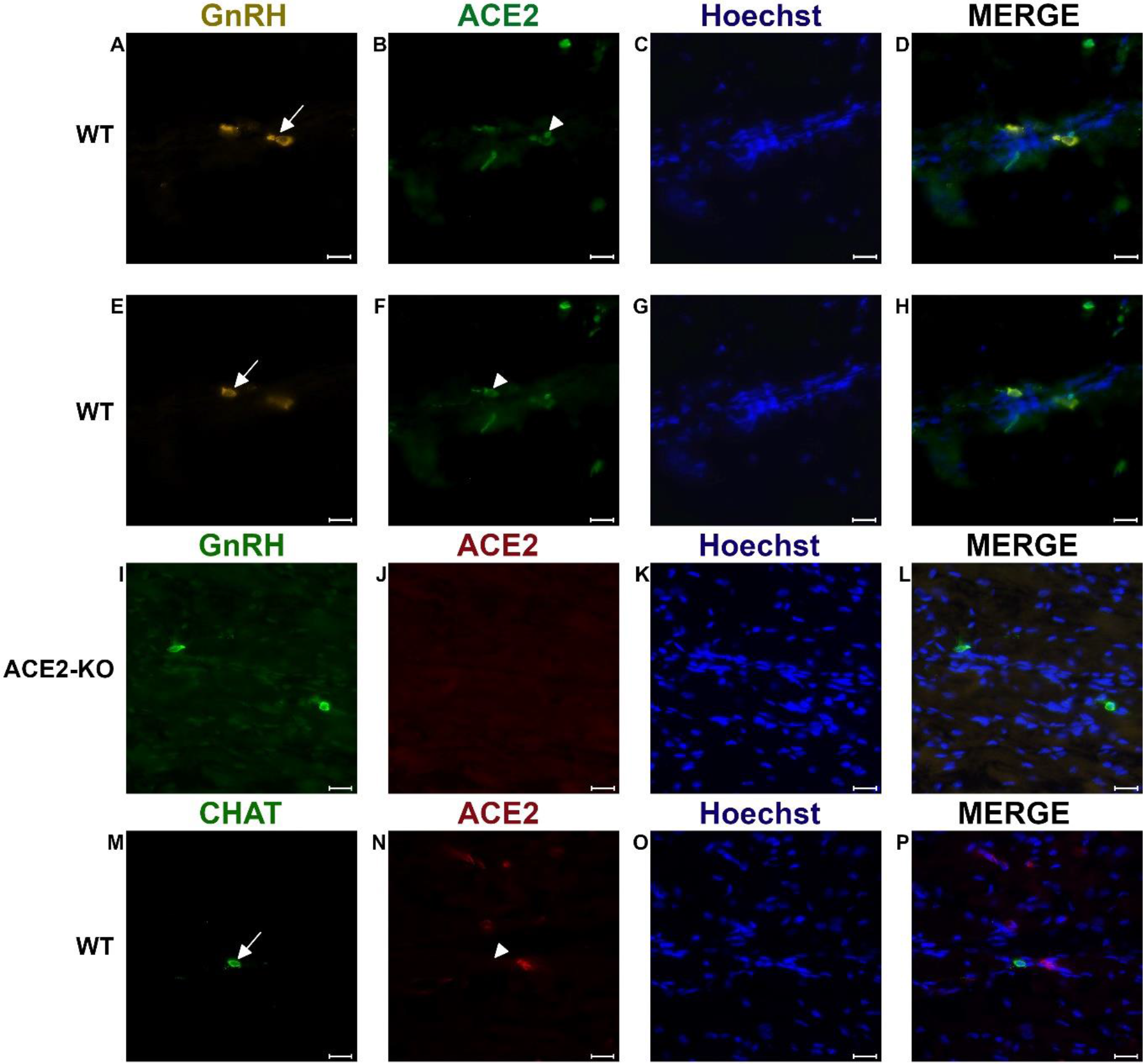
Examples of double immunofluorescent labeling for nervus terminalis neuronal markers GnRH (**A, E**) or CHAT (**M**) and ACE2 (**B, F, J, N**) in the medial region adjacent to the olfactory bulbs as indicated in Fig. 1. Panels **A-D** and **E-H** show slightly different focal planes to demonstrate the morphology of the two or three different neurons. Nuclei are stained with Hoechst 33258 (**C, G, K, O**). Merged images are shown in the last column (**D, H, L, P**). The neuronal somas labeled with GnRH (**A, E**) are co-labeled with ACE2 (**B, F**) as shown after merging (**D, H**). GnRH positive cells in the ACE2 knock-out mouse (**I**) are not labeled with ACE2 (**J**). The majority of cholinergic neurons are not labeled with ACE2 (**M, N**), as quantified in Fig. 3A. Control sections probed without primary antibodies or with control rabbit IgG had no detectable signal (not shown). Arrows and triangles indicate double-labeled neurons or lack thereof. Scale bars: 20 μm.

### Most GnRH^+^ nervus terminalis neurons express ACE2

To determine whether some neurons of the nervus terminalis contained both GnRH and ACE2, and to estimate the number of such neurons, we performed double immunolabeling experiments, and single- and double-labeled cells were counted on 15-20 sections from five different animals. The analyzed olfactory epithelium and olfactory bulb region and section’s cutting plane are as indicated in Fig. 1. Out of 119 GnRH+ neurons, 107 (89.9%) were double-labeled for ACE2 (Fig. 3A). The intensity (0-250 scale) of ACE2 immunolabel in nervus terminalis neurons (range of 63 to 114, mean=86) was comparable to the intensity of ACE2 immunolabel (range of 70 to 141, mean=108) that we have previously demonstrated for sustentacular cells in the olfactory epithelium in some of the same tissue sections, and with the same protocol (Bilinska et al., 2020). Controls included omission of the primary antibody (not shown) and double immunofluorescent reactions performed using cryosections derived from an ACE2 knockout mouse as shown in Fig. 2 (I-L). The GnRH-positive neurons were never labeled for olfactory marker protein (OMP), a marker for mature olfactory receptor neurons (Fig. S1, Supplementary Material). The total number of GnRH+ nervus terminalis neurons per mouse was estimated to be approximately 125, which is very similar to a previous serial section analysis in hamster (about 130-140 GnRH+ neurons, Wirsig and Leonard, 1986).

**Fig. 3A-F.**
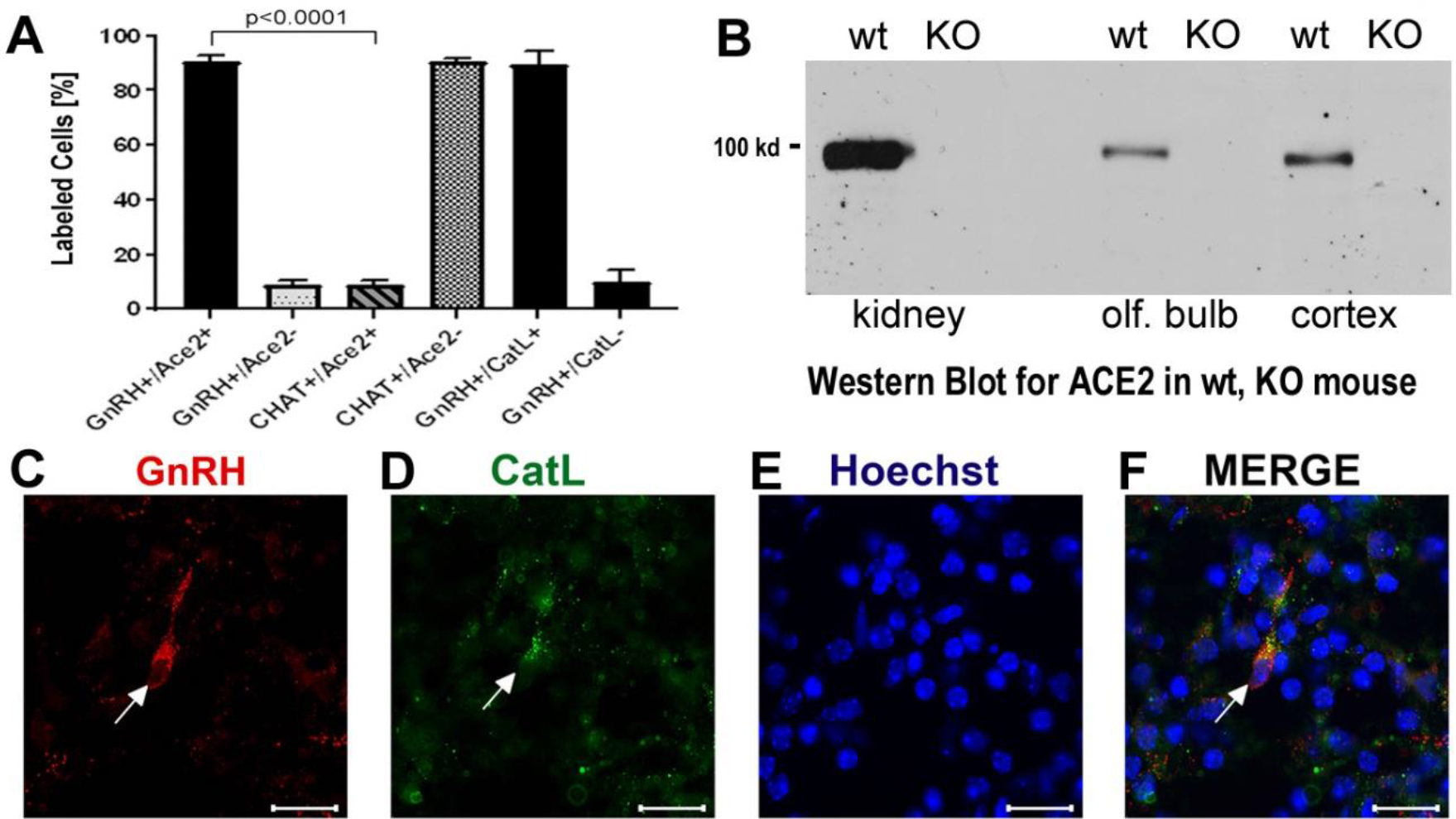
Quantification of neurons labeled with nervus terminalis markers, virus entry proteins, and verification of the specificity of the ACE2 antibody. **A.** The large majority of GnRH-positive neurons is also ACE2-positive. In contrast, the majority of CHAT-positive (cholinergic) nervus terminalis neurons lack ACE2-expression. The total number of counted GnRH-positive or CHAT-positive neurons was set at 100%. Error bars represent ± SEM. A t-test shows that the colocalization difference between GnRH- and CHAT-positive nervus terminalis neurons is significant at p<0.0001. For further details, see Table S2. **B.** Western blot of ACE2 in wildtype (wt) mice and in ACE2 knock-out (KO) mice. The first two lanes (kidney) were loaded with 25 μg total protein, the lanes for olfactory bulb and cerebral cortex were loaded with 60 μg total protein, and probed with the R&D ACE2 antibody. No ACE2 protein was detectable in the ACE2 KO mice, proving that the antibody indeed recognizes ACE2. **C-F.** Example of GnRH-positive nervus terminalis neurons which are also cathepsin L-positive. **C.** One GnRH-labeled neuron is marked with a white arrow. **D.** The same neuron is labeled with the cathepsin L antibody (CatL, white arrow). **E.** The cell nuclei are stained with Hoechst nuclear dye. **F.** The three images are merged to show co-localization in the neuron indicated with the white arrow. All scale bars are 20 μm.

### A small fraction of CHAT^+^ nervus terminalis neurons express ACE2

The nervus terminalis complex is comprised of several distinct heterogenous populations of neurons. In addition to GnRH neurons which form the major fraction of nervus terminalis neurons, the second largest nervus terminalis subpopulation are cholinergic neurons that can be identified by the presence of choline acetyltransferase (CHAT) or cholinesterase (Wirsig and Leonard, 1986; Demski, 1993; von Bartheld, 2004). Therefore, CHAT neurons were also double-labeled with ACE2 and the fraction of CHAT-positive and ACE2-positive neurons was estimated out of a total of 52 CHAT-positive neurons in three animals. In contrast to GnRH+/ACE2+ cells, only a minor fraction (9.4%) of CHAT-positive neurons were labeled with ACE2 which indicates that relatively few cholinergic nervus terminalis neurons express ACE2 protein (Fig. 3A, Table S2). Most of the CHAT-positive and also ACE2-positive nervus terminalis neurons were fusiform and unipolar in shape. Control experiments included omission of primary antibody (not shown) and double immunofluorescent reactions performed using cryosections derived from ACE2 knockout mouse (see below). The total number of CHAT-positive nervus terminalis neurons per mouse was estimated to be approximately 60-70. This is less than the 130-140 acetylcholinesterase containing nervus terminalis neurons in hamster (Wirsig and Leonard, 1986), but it is known that only a fraction of neurons containing acetylcholinesterase actually are cholinergic (Schwanzel-Fukuda et al., 1986). The difference in colocalization of ACE2 between the two transmitter phenotypes of nervus terminalis neurons was highly significant, indicating that ACE2 expression is not randomly distributed in the nervus terminalis system.

### Proof of the specificity of the ACE2 antibody: Western blots from ACE2 −/− mice

Immunolabeling experiments did not reveal any signal beyond background when the primary antibodies were omitted. For more precise visualization of background ACE2 staining, tissue derived from ACE2 knock-out mouse was used. In Western blots, the ACE2-specific band was absent in total protein extract obtained from ACE2 −/− animals (Fig. 3B). Therefore, sections from ACE2 −/− mice were also used for double immunolabeling experiments with ACE2 antibody and results showed that, as expected, GnRH-positive neurons were negative for ACE2 in tissue sections from the knock-out animals (Fig. 2 I-L).

### The protease TMPRSS2 is not or minimally expressed in nervus terminalis neurons

TMPRSS2 is one of several proteases that cleave the spike protein, allowing SARS-CoV-2 to enter the host cell (Shang et al., 2020). Our attempts to double-label nervus terminalis neurons using three different anti-TMPRSS2 antibodies (Table S1) resulted in no specific label above background in GnRH-positive nervus terminalis neurons (Supplemental Material, Fig. S1E). Levels of TMPRSS2 expression in these neurons may be too low for detection, or TMPRSS2 is not expressed by either the GnRH- or CHAT-expressing nervus terminalis neurons.

### The proteases cathepsins B and L are expressed in nervus terminalis neurons

It is known that endolysosomal proteases cathepsin B and L can also enhance entry of SARS-CoV-2 to host cells (Gomes et al., 2020; Bollavaram et al., 2021; Zhao et al., 2021). To determine whether some neurons of the nervus terminalis contained both GnRH and cathepsin B or L, and to estimate the number of such cells, we performed double immunolabeling experiments, and single- and double-labeled cells were counted on 15-20 sections from three different animals. The analyzed olfactory epithelium and olfactory bulb region and section’s cutting plane are as indicated in Fig. 1. After surveying a total of 51 GnRH positive neurons, we found that 90.0% of them were double-labeled for cathepsin L (Cath L, Fig. 3A). A similar trend was observed for cathepsin B with a large proportion of GnRH-positive neurons co-labeled with the two antibodies (data not shown).

## DISCUSSION

Our experiments confirmed the locations and approximate numbers of GnRH-positive neurons of the nervus terminalis in rodents (Schwanzel-Fukuda et al., 1986; Wirsig and Leonard, 1986; Schwanzel-Fukuda et al., 1987). In mouse, we found that the number of CHAT-positive neurons was about half of the number of GnRH-positive neurons. Interestingly, the large majority of GnRH-positive neurons expressed ACE2 and cathepsin B or L, while only a small fraction of CHAT-positive neurons co-localized ACE2. Previous studies have suggested that a larger fraction of the CHAT-positive neurons were multipolar and possibly associated with an autonomic function, while GnRH-positive neurons are thought to be sensory and/or may have neurosecretory functions (Wirsig and Leonard, 1986; Schwanzel-Fukuda et al., 1986).

Mice have been most often used as model systems for ACE2 expression, for localization of SARS-CoV-2 in the olfactory epithelium, and to study neuro-invasion of the brain along the olfactory route (Butowt and von Bartheld, 2020; Cooper et al., 2020; Rathnasinghe et al., 2020). Mice have the advantage that a large number of mutants are available (Butowt and von Bartheld, 2020), but they normally express an ACE2 version that binds SARS-CoV-2 with low affinity (Damas et al., 2020). Therefore, to study SARS-CoV-2 infection in mice, a mouse-adapted virus has to be used (Leist et al., 2020), or mice have to be engineered to express human ACE2 (Butowt et al., 2021).

SARS-CoV-2 uses ACE2 to bind to host cells, and then the spike protein is cleaved by surface or endosomal proteases to facilitate virus entry. One of the more common proteases to facilitate SARS-CoV-2 entry is TMPRSS2 (Shang et al., 2020). TMPRSS2 is minimally expressed in adult neurons (Paoloni-Giacobino et al., 1997), including GnRH-expressing neurons in the brain (Ubuka et al., 2018). Similarly, TMPRSS2 was minimally or not at all expressed in nervus terminalis neurons. TMPRSS2 is only one of several proteases that are known to cleave the spike protein of SARS-CoV-2. Additional proteases that can facilitate SARS-CoV-2 entry into host cells in the absence of TMPRSS2 include cathepsins B and L (Butowt et al., 2020; Gomes et al., 2020; Bollavaram et al., 2021; Zhao et al., 2021), and also furin. Pre-activation by furin enhances viral entry in cells that lack TMPRSS2 or cathepsins, or have low levels of expression of these proteases (Shang et al., 2020). Furin is a ubiquitously expressed protease with a fundamental role in maturation of proteins in the secretory pathway; it is expressed in virtually all cells in rodent brain albeit at different levels (Day et al., 1993). Furin is known to be expressed in Bowman gland cells (Ueha et al., 2021), and therefore would not be a limiting factor for SARS-CoV-2 entry into innervating nervus terminalis neurons. Accordingly, lack of TMPRSS2 in nervus terminalis neurons does not negate the possibility of SARS-CoV-2 infecting these neurons after using alternative proteases to gain entry.

Our finding of expression of ACE2 and cathepsins B and L in the large majority of GnRH-expressing nervus terminalis neurons suggests that this cranial nerve is a more plausible conduit for brain infection than the olfactory neurons that entirely or for the most part lack ACE2 expression (Butowt and von Bartheld, 2020; Cooper et al., 2020; Klingenstein et al., 2021). The nervus terminalis neurons may obtain the SARS-CoV-2 directly from infected cells in Bowman’s glands, or through free nerve endings within the olfactory epithelium, many parts of which degenerate when sustentacular cells are infected by SARS-CoV-2 (Bryche et al., 2020). The lack of ACE2 in olfactory receptor neurons (except for those in the Grueneberg ganglion, Brechbühl et al., 2021) appears to be an effective barrier to virus transfer along the olfactory nerve (Butowt et al., 2021). The nervus terminalis, however, has multiple venues to become infected by the virus, as illustrated in Fig. 4.

**Fig. 4.**
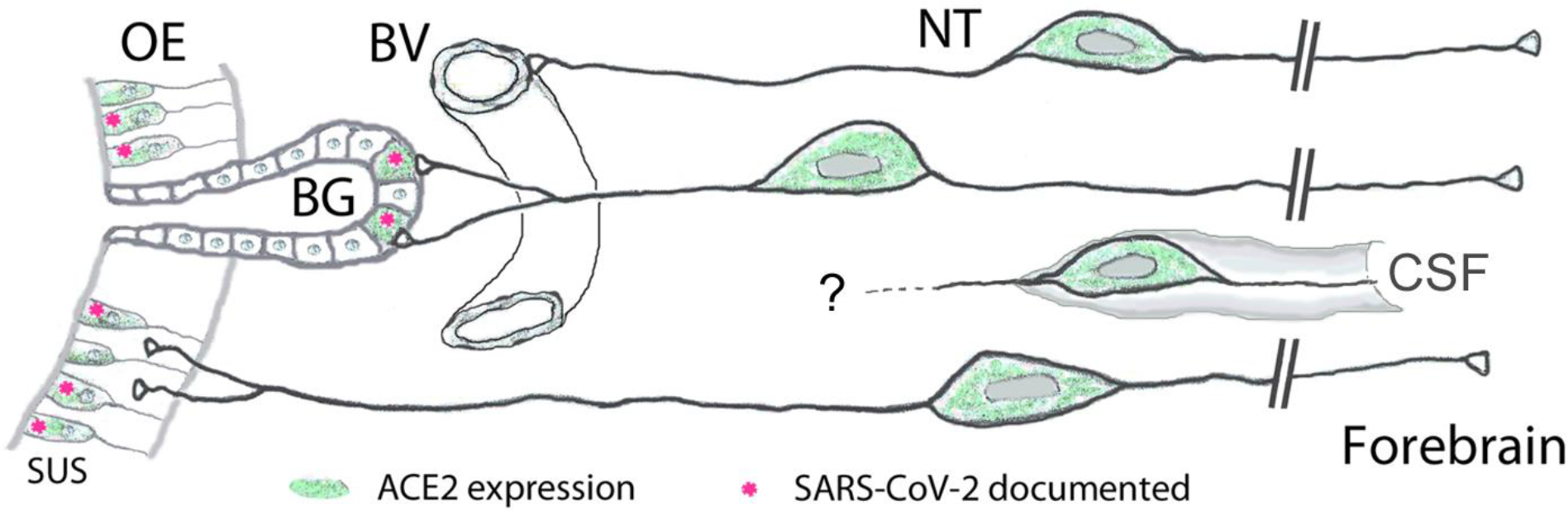
Peripheral projections of nervus terminalis (NT) neurons and their presumptive relationship with ACE2-expressing neurons in the olfactory epithelium and known SARS-CoV-2 infection. NT neurons innervate blood vessels (BV), Bowman gland (BG) cells, the olfactory epithelium (OE), and contact cerebrospinal fluid (CSF) spaces. Peripheral projections of NT neurons according to Larsell (1950), CSF contacts according to Jennes (1987). Cells expressing ACE2 are indicated in green, including sustentacular cells (SUS) and BG cells. Both of these cell types have been shown to express ACE2 (Bilinska et al., 2020; Brann et al., 2020; Chen et al., 2020; Ye et al., 2020; Zhang et al., 2020; Klingenstein et al., 2021). Cell types that have been documented to be infected by SARS-CoV-2 are indicated with pink asterisks. SARS-CoV-2 localization in SUS cells according to Bryche et al., 2020; Leist et al., 2020; Ye et al., 2020; Zhang et al., 2020; Zheng et al., 2020; de Melo et al., 2021, and in BG cells according to Bryche et al., 2020; Leist et al., 2020; Ye et al., 2020. BG cells furthermore express the protease furin (Ueha et al., 2021) which may facilitate virus entry into those nervus terminalis neurons which innervate BG cells.

Another important aspect is that the timeline of appearance of SARS-CoV-2 in the brain fits the nervus terminalis projections, with an explosive appearance of the virus in the forebrain in some mouse models (Winkler et al., 2020; Zheng et al., 2020; Zhou et al., 2020; Carossino et al., 2021), rather than a gradual transfer along the olfactory projections as would be expected from a virus that gains access to the brain via olfactory projections (Barnett and Perlman, 1993). The nervus terminalis has direct projections into the forebrain, reaching as far caudal as the hypothalamus (von Bartheld, 2004), and if the virus indeed infects these neurons, this could explain why the virus reaches the brain and cerebrospinal fluid (CSF) spaces much faster than seems possible via “neuron hopping” along olfactory projections. Most of the virus-containing axons in the olfactory nerve demonstrated by de Melo et al. (2021) do not express olfactory marker protein, suggesting that they are not axons belonging to olfactory receptor neurons, and therefore may be nervus terminalis axons which also project along the olfactory nerve (Larsell, 1950).

On a comparative note, since dolphins and whales have a much larger number of nervus terminalis neurons than any other vertebrates (Oelschläger et al., 1987), and these marine mammals express ACE2 that is highly susceptible to SARS-CoV-2 binding (Damas et al., 2020), our finding of ACE2 in nervus terminalis neurons suggests that these animals may be more vulnerable to brain infection via the nervus terminalis – even in the absence of an olfactory system.

In humans, the number of nervus terminalis neurons is relatively small (a few hundred to a few thousand neurons depending on age, Brookover, 1917; Larsell, 1950; Jin et al., 2019). However, it is possible that such a relatively small number is sufficient to mediate viral infection. The nervus terminalis directly innervates secretory cells of the Bowman’s glands (Larsell, 1950) that are known to express ACE2 (Brann et al., 2020; Chen et al., 2020; Cooper et al., 2020; Ye et al., 2020; Zhang et al., 2020; Klingenstein et al., 2021) and readily become infected with SARS-CoV-2 (Ye et al., 2020; Leist et al., 2020; Meinhardt et al., 2020; Zhang et al., 2020; Zheng et al., 2020) (Fig. 4). In addition, the nervus terminalis has many free nerve endings within the olfactory epithelium (Larsell, 1950) – an epithelium that is heavily damaged when ACE2-expressing sustentacular cells become infected and degenerate (Bryche et al., 2020). Finally, a major component of the nervus terminalis innervates blood vessels below the olfactory epithelium and projects via cerebrospinal fluid (CSF)-containing spaces (Larsell, 1950; Jennes, 1987). Some nervus terminalis neurons have direct projections to the hypothalamus (Pearson, 1941; Larsell, 1950; von Bartheld, 2004), a brain region that may serve as a hub for virus spread throughout the brain (Nampoothiri et al., 2020; Zheng et al., 2020).

Another argument to consider the nervus terminalis as an alternative to the olfactory route is that neuro-invasion in most animal models is highly variable, even in the same species and transgenic model (Jiang et al., 2020; Oladunni et al., 2020; Rathnasinghe et al., 2020; Winkler et al., 2020; Ye et al., 2020; Zheng et al., 2020; Zhou et al., 2020), and this is in contrast to the olfactory system that is consistent in terms of numbers of neurons, gene expression and projections. The nervus terminalis, on the other hand, is known for its large variability between individuals of the same species or even when comparing the right side with the left side of the same individual (Larsell, 1918; Jin et al., 2019). Such numerical differences can approach or even exceed an entire order of magnitude (Schwanzel-Fukuda et al., 1987; Jin et al., 2019) – and thus may explain the reported large variability in neuro-invasion (Butowt et al., 2021). Taken together, nervus terminalis neurons, for the above reasons, should be considered as a plausible alternative to the olfactory projections for neuro-invasion of SARS-CoV-2 from the nose to the brain in COVID-19. Whether the virus can indeed infect nervus terminalis neurons cannot be deduced by protein expression, but has to be determined by experiments using infectious SARS-CoV-2 after nasal inoculation in experimental animal models, or by demonstration of SARS-CoV-2 in nervus terminalis neurons in tissues from patients with COVID-19.

## ACKNOWLEDGMENTS

Supported by the “Excellence Initiative-Research University" programme at the Nicolaus Copernicus University (R.B.), and grant GM103554 from the National Institutes of Health (C.S.v.B.).

## CONTRIBUTION TO THE FIELD

The new coronavirus responsible for the COVID-19 pandemic can infect the brain in humans and in some animal models. It is currently not known how this virus infects the brain. Many researchers believe that the virus enters the brain by using a route along the olfactory nerve. However, the olfactory neurons in the nose do not express the obligatory virus entry receptors, and the timing of arrival and transfer of the virus in brain targets is inconsistent with a neuronal transfer along olfactory projections. Here we show that an alternative route for the new coronavirus to infect the brain is more plausible. We show that many nervus terminalis neurons express the spike-binding virus entry protein. Since these neurons have direct synaptic contact with cells known to become infected in the nose, and have direct projections to various targets in the forebrain, the nervus terminalis neurons may provide an alternative route for the new coronavirus to gain access from the nose to the brain. The main contribution of our paper is to show that a plausible alternative route to brain infection exists that needs to be pursued by examination of human tissues and by testing in appropriate animal models whether the new coronavirus indeed utilizes this pathway.

## SUPPLEMENTARY MATERIAL

### SUPPLEMENTAL TABLES

**Table S1.** Primary and secondary antibodies used in this study.

| Primary antibodies | Company | Catalog # | Type |
| --- | --- | --- | --- |
| ACE2 | R&D Systems | AF3437 | goat polyclonal |
| ACE2 | ABclonal | A4612 | rabbit monoclonal |
| Cathepsin B | R&D Systems | AF965 | goat polyclonal |
| Cathepsin L | R&D Systems | AF1515 | goat polyclonal |
| CHAT | Proteintech | 24418-1-AP | rabbit polyclonal |
| GnRH1 | Proteintech | 26950-1-AP | rabbit polyclonal |
| OMP | WAKO | 544-10001 | goat polyclonal |
| TMPRSS2 | Novus Biologicals | NBP3-00492 | rabbit monoclonal |
| TMPRSS2 | Abcam | ab242384 | rabbit monoclonal |
| TMPRSS2 | St John's | STJ11102428 | rabbit monoclonal |
| Secondary antibodies |  |  | Fluorescent conjugate |
| donkey anti-rabbit | Abcam | ab15006 | Alexa Fluor 488 |
| donkey anti-goat | Abcam | ab150136 | Alexa Fluor 594 |
| horse anti-rabbit | Vector | DI-1094 | Alexa Fluor 594 |
| horse anti-goat | Vector | DI-3088 | Alexa Fluor 488 |

Abbreviations: ACE2, angiotensin converting enzyme 2; CHAT, choline acetyltransferase; GnRH1, gonadotropin releasing hormone 1; OMP, olfactory marker protein; TMPRSS2, transmembrane protease serine 2.

**TABLE S2.** Numbers of animals, sections, and numbers of neurons double-labeled for nervus terminalis markers (GnRH, CHAT) and ACE2.

|  | # of sections | # of GnRH neurons | # of GnRH and ACE2 | % |
| --- | --- | --- | --- | --- |
| **GnRH and ACE2** |  |  |  |  |
| Mouse 1 | 16 | 25 | 23 | 92.0 |
| Mouse 2 | 21 | 30 | 27 | 90.0 |
| Mouse 3 | 15 | 21 | 19 | 90.5 |
| Mouse 4 | 11 | 10 | 8 | 80.0 |
| Mouse 5 | 22 | 33 | 30 | 90.9 |
| **Sum/mean** | **85** | **119** | **107** | **89.9** |
| **CHAT and ACE2** |  |  |  |  |
| Mouse 1 | 21 | 20 | 2 | 10.0 |
| Mouse 2 | 15 | 18 | 2 | 11.1 |
| Mouse 3 | 18 | 14 | 1 | 7.1 |
| **Sum/mean** | **54** | **52** | **5** | **9.4** |
| **t-test** |  |  |  | **p<0.0001** |

### SUPPLEMENTAL FIGURES

**Fig. S1.**
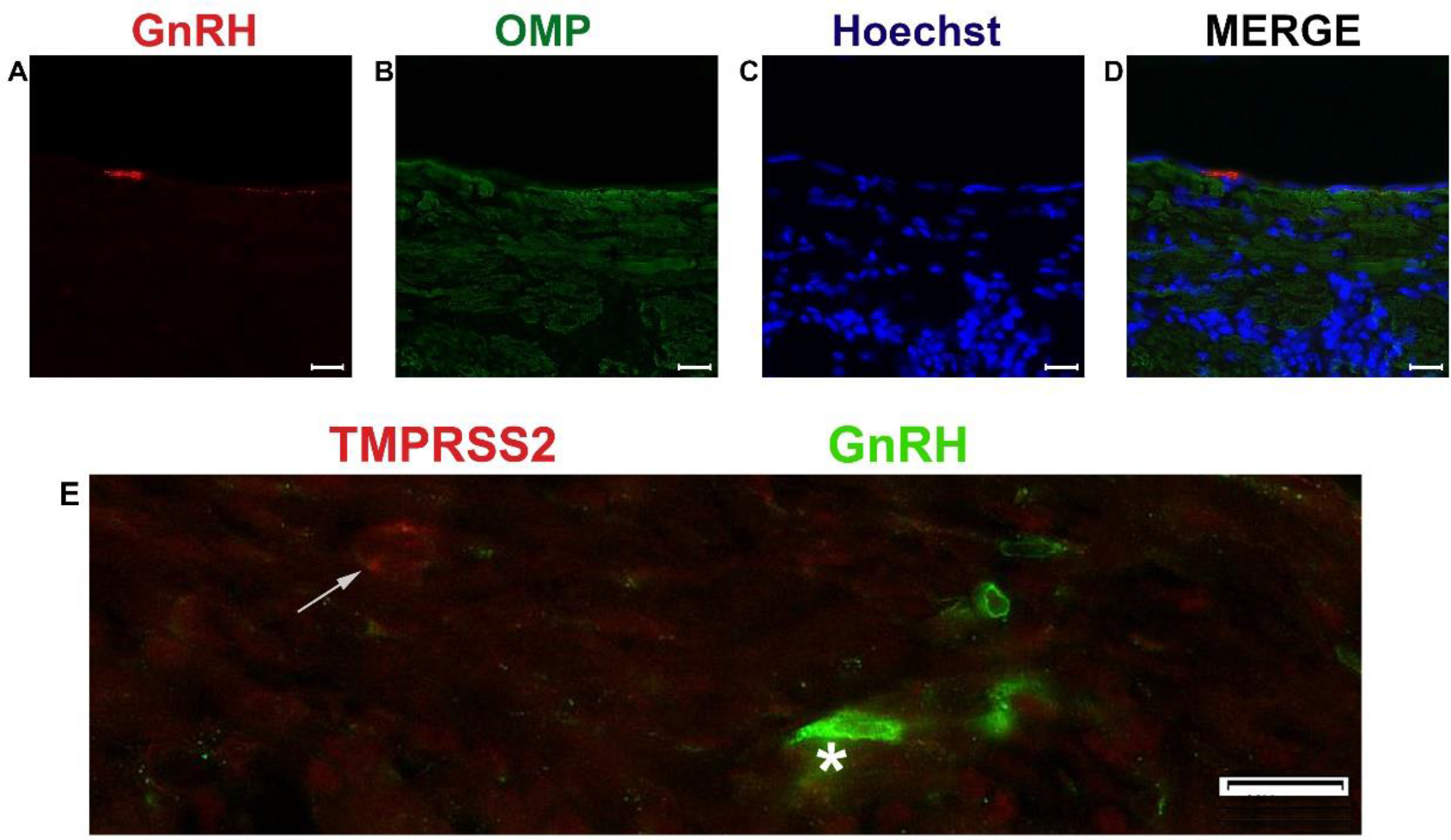
Examples of double immunofluorescent labeling of nervus terminalis neurons with the markers GnRH and olfactory marker protein (OMP) (**A-D**), and GnRH and TMPRSS2 (**E**). Label for GnRH (**A**) and OMP (**B**) in the medial region adjacent to the olfactory bulbs as indicated in Fig. 1. Nuclei are stained with Hoechst 33258 (**C**) and the merged image is shown in (**D**). GnRH-labeled cells were never labeled for OMP in this region. (**E**) GnRH-labeled nervus terminalis neurons (one neuron indicated with the white asterisk) did not co-localize with cells positive for TMPRSS2 (white arrow). Scale bars: 20 μm.

## REFERENCES

Barnett, E.M., and Perlman, S. (1993). The olfactory nerve and not the trigeminal nerve is the major site of CNS entry for mouse hepatitis virus, strain JHM. Virology. 194(1), 185–191.

Barnett, E.M., Evans, G.D., Sun, N., Perlman, S., and Cassell, M.D. (1995). Anterograde tracing of trigeminal afferent pathways from the murine tooth pulp to cortex using herpes simplex virus type 1. J Neurosci. (4), 2972–2984.

Bilinska, K., Jakubowska, P., Von Bartheld, C.S., and Butowt, R. (2020). Expression of the SARS-CoV-2 Entry Proteins, ACE2 and TMPRSS2, in Cells of the Olfactory Epithelium: Identification of Cell Types and Trends with Age. ACS Chem Neurosci. 11(11), 1555–1562.

Bollavaram, K., Leeman, T. H., Lee, M. W., Kulkarni, A., Upshaw, S. G., Yang, J., et al. (2021). Multiple sites on SARS-CoV-2 spike protein are susceptible to proteolysis by cathepsins B, K, L, S, and V. Protein Sci. 2021 Jun;30(6):1131–1143.

Bougakov, D., Podell, K., and Goldberg, E. (2020). Multiple Neuroinvasive Pathways in COVID-19. Mol Neurobiol. 29,1–12.

Briguglio, M., Bona, A., Porta, M., Dell’Osso, B., Pregliasco, F.E., and Banfi, G. (2020). Disentangling the hypothesis of host dysosmia and SARS-CoV-2: The bait symptom that hides neglected neurophysiological routes. Front Physiol. 11, 671.

Brann, D.H., Tsukahara, T., Weinreb, C., Lipovsek, M., Van den Berge, K., Gong, B., et al. (2020). Non-neuronal expression of SARS-CoV-2 entry genes in the olfactory system suggests mechanisms underlying COVID-19-associated anosmia. Sci Adv. 6(31):eabc5801.

Brechbühl, J., Wood, D., Bouteiller, S., Lopes, A.C., Verdumo, C., and Broillet, M.-C. (2021). Age-dependent appearance of SARS-CoV-2 entry cells in mouse chemosensory systems reflects COVID-19 anosmia and ageusia symptoms. bioRxiv [Preprint] March 29, 2021 doi: https://doi.org/10.1101/2021.03.29.437530.

Brookover, C. (1917). The peripheral distribution of the nervus terminalis in an infant. J Comp Neurol. 28, 349–360.

Bryche, B., St Albin, A., Murri, S., Lacôte, S., Pulido, C., Ar Gouilh, M., et al. (2020). Massive transient damage of the olfactory epithelium associated with infection of sustentacular cells by SARS-CoV-2 in golden Syrian hamsters. Brain Behav Immun. 89, 579–586.

Burks, S.M., Rosas-Hernandez, H., Alenjandro Ramirez-Lee, M., Cuevas, E., and Talpos, J.C. (2021). Can SARS-CoV-2 infect the central nervous system via the olfactory bulb or the blood-brain barrier? Brain Behav Immun. 20, 32489–32492.

Butowt, R., and Bilinska, K. (2020). SARS-CoV-2: Olfaction, Brain Infection, and the Urgent Need for Clinical Samples Allowing Earlier Virus Detection. ACS Chem Neurosci. 11(9),1200–1203.

Butowt, R., and von Bartheld, C.S. (2020). Anosmia in COVID-19: Underlying Mechanisms and Assessment of an Olfactory Route to Brain Infection. Neuroscientist. doi: 10.1177/1073858420956905. Epub ahead of print.

Butowt, R., Pyrc, K., and von Bartheld, C.S. (2020). Battle at the entrance gate: CIITA as a weapon to prevent the internalization of SARS-CoV-2 and Ebola viruses. Signal Transduct Target Ther. Nov 24;5(1):278.

Butowt, R., Meunier, N., Bryche, B., and von Bartheld, C.S. (2021). The olfactory nerve is not a likely route to brain infection in COVID-19: a critical review of data from humans and animal models. Acta Neuropathol, 1–14. Advance online publication. https://doi.org/10.1007/s00401-021-02314-2.

Carossino, M., Montanaro, P., O’Connell, A., Kenney, D., Gertje, H., Grosz, K.A., et al. (2021). Fatal neuroinvasion of SARS-CoV-2 in K18-hACE2 mice is partially dependent on hACE2 expression. bioRxiv [Preprint] doi: 10.1101/2021.01.13.425144. (accessed on May 30, 2021).

Chen, M., Shen, W., Rowan, N. R., Kulaga, H., Hillel, A., Ramanathan, M., Jr, et al. (2020). Elevated ACE-2 expression in the olfactory neuroepithelium: implications for anosmia and upper respiratory SARS-CoV-2 entry and replication. Eur Resp J. 56(3), 2001948.

Cooper, K.W., Brann, D.H., Farruggia, M.C., Bhutani, S., Pellegrino, R., Tsukahara, T., et al. (2020). COVID-19 and the Chemical Senses: Supporting Players Take Center Stage. Neuron. 107(2), 219–233.

Damas, J., Hughes, G.M., Keough, K.C., Painter, C.A., Persky, N.S., Corbo, M., et al. (2020). Broad host range of SARS-CoV-2 predicted by comparative and structural analysis of ACE2 in vertebrates. Proc Natl Acad Sci U S A. 117(36), 22311–22322.

Day, R., Schafer, M.K., Cullinan, W.E., Watson. S.J., Chrétien. M., Seidah, N.G. (1993). Region specific expression of furin mRNA in the rat brain. Neurosci Lett. 149(1), 27–30.

de Melo, G.D., Lazarini, F., Levallois, S., Hautefort, C., Michel, V., Larrous, F., et al. (2021). COVID-19-related anosmia is associated with viral persistence and inflammation in human olfactory epithelium and brain infection in hamsters. Sci Transl Med, eabf8396. E-pub. https://doi.org/10.1126/scitranslmed.abf8396.

Demski, L.S. (1993). Terminal nerve complex. Acta Anat (Basel). 148(2-3), 81–95.

Dubé, M., Le Coupanec, A., Wong, A.H.M., Rini, J.M., Desforges, M., and Talbot, P.J. (2018). Axonal Transport Enables Neuron-to-Neuron Propagation of Human Coronavirus OC43. J Virol. 92(17), 404–418.

Ellul, M.A., Benjamin, L., Singh, B., Lant, S., Michael, B.D., Easton, A., et al. (2020). Neurological associations of COVID-19. Lancet Neurol. 19(9), 767–783.

Gomes, C. P., Fernandes, D. E., Casimiro, F., da Mata, G. F., Passos, M. T., Varela, P., et al. (2020). Cathepsin L in COVID-19: From Pharmacological Evidences to Genetics. Front Cell Infect Microbiol. Dec 8;10:589505.

Jennes, L. (1987). The nervus terminalis in the mouse: light and electron microscopic immunocytochemical studies. Ann N Y Acad Sci. 519,165–173.

Jiang, R. D., Liu, M. Q., Chen, Y., Shan, C., Zhou, Y. W., Shen, X. R., et al. (2020). Pathogenesis of SARS-CoV-2 in Transgenic Mice Expressing Human Angiotensin-Converting Enzyme 2. Cell. 182(1), 50–58.e8.

Jin, Z.W., Cho, K.H., Shibata, S., Yamamoto, M., Murakami, G., and Rodríguez-Vázquez, J.F. (2019). Nervus terminalis and nerves to the vomeronasal organ: a study using human fetal specimens. Anat Cell Biol. 52(3), 278–285.

Kim, K.H., Patel, L., Tobet, S.A., King, J.C., Rubin, B.S., and Stopa, E.G. (1999). Gonadotropin-releasing hormone immunoreactivity in the adult and fetal human olfactory system. Brain Res. 826(2), 220–229.

Klingenstein, M., Klingenstein, S., Neckel, P.H., Mack, A.F., Wagner, A.P, Kleger, A., et al. (2021). Evidence of SARS-CoV2 Entry Protein ACE2 in the Human Nose and Olfactory Bulb. Cells Tissues Organs 209(4-6), 155–164.

Larsell, O. (1918). Nervus terminalis: mammals. J Comp Neurol. 30, 3–68.

Larsell, O. (1950). The nervus terminalis. Ann Otol Rhinol Laryngol. 59(2), 414–438.

Leist, S.R., Dinnon, K.H., Schäfer, A., Tse, L.V., Okuda, K., Hou, Y.J., et al. (2020). A Mouse-Adapted SARS-CoV-2 Induces Acute Lung Injury and Mortality in Standard Laboratory Mice. Cell. 183 (4),1070–1085.

Li, Z., Liu, T., Yang, N., Han, D., Mi, X., Li, Y., et al. (2020). Neurological manifestations of patients with COVID-19: potential routes of SARS-CoV-2 neuroinvasion from the periphery to the brain. Front Med. 14(5), 533–541.

Liang, F. (2020). Sustentacular Cell Enwrapment of Olfactory Receptor Neuronal Dendrites: An Update. Genes (Basel). 11(5), 493.

Masre, S.F., Jufri, N.F., Ibrahim, F.W., and Abdul Raub, S.H. (2020). Classical and alternative receptors for SARS-CoV-2 therapeutic strategy. Rev Med Virol. e2207. Advance online publication. https://doi.org/10.1002/rmv.2207.

Matschke, J., Lütgehetmann, M., Hagel, C., Sperhake, J.P., Schröder, A.S., Edler, C., et al. (2020). Neuropathology of patients with COVID-19 in Germany: a post-mortem case series. Lancet Neurol. 19(11), 919–929.

Meinhardt, J., Radke, J., Dittmayer, C., Franz, J., Thomas, C., Mothes, R., et al. (2021). Olfactory transmucosal SARS-CoV-2 invasion as a port of central nervous system entry in individuals with COVID-19. Nat Neurosci. 24(2), 168–175.

Mukerji, S.S., and Solomon, I.H. (2021). What can we learn from brain autopsy in COVID-19? Neurosci Lett. 742:135528. doi: 10.1016/j.neulet.2020.135528.

Nampoothiri, S., Sauve, S., Ternier, G., Fernandois, D., Coelho, C., Imbernon, M., et al. (2020). The hypothalamus as a hub for putative SARS-CoV-2 brain infection. bioRxiv [Preprint] doi: https://doi.org/10.1101/2020.06.08.139329. (accessed on May 30, 2021)

Natoli, S., Oliveira, V., Calabresi, P., Maia, L.F., and Pisani, A. (2020). Does SARS-Cov-2 invade the brain? Translational lessons from animal models. Eur J Neurol. 9,1764–1773.

Netland, J., Meyerholz, D.K., Moore, S., Cassell, M., and Perlman, S. (2008). Severe acute respiratory syndrome coronavirus infection causes neuronal death in the absence of encephalitis in mice transgenic for human ACE2. J Virol. 82(15), 7264–7275.

Oelschläger, H.A., Buhl, E.H., and Dann, J.F. (1987). Development of the nervus terminalis in mammals including toothed whales and humans. Ann N Y Acad Sci. 519, 447–464.

Oladunni, F. S., Park, J. G., Pino, P. A., Gonzalez, O., Akhter, A., Allué-Guardia, A., et al. (2020). Lethality of SARS-CoV-2 infection in K18 human angiotensin-converting enzyme 2 transgenic mice. Nat Commun. 11(1), 6122.

Paoloni-Giacobino, A., Chen, H., Peitsch, M. C., Rossier, C., and Antonarakis, S.E. (1997). Cloning of the TMPRSS2 gene, which encodes a novel serine protease with transmembrane, LDLRA, and SRCR domains and maps to 21q22.3. Genomics, 44(3), 309–320.

Pearson, A.A. (1941). The development of the nervus terminalis in man. J Comp Neurol. 75, 39–66.

Rathnasinghe, R., Strohmeier, S., Amanat, F., Gillespie, V.L., Krammer, F., García-Sastre, A., et al. (2020). Comparison of transgenic and adenovirus hACE2 mouse models for SARS-CoV-2 infection. Emerg Microbes Infect. 1, 2433–2445.

Ridgway, S.H., Demski, L.S., Bullock, T.H., and Schwanzel-Fukuda, M. (1987). The terminal nerve in odontocete cetaceans. Ann N Y Acad Sci. 519, 201–212.

Schwanzel-Fukuda, M., Morrell, J. I., and Pfaff, D. W. (1986). Localization of choline acetyltransferase and vasoactive intestinal polypeptide-like immunoreactivity in the nervus terminalis of the fetal and neonatal rat. Peptides. 7(5), 899–906.

Schwanzel-Fukuda, M., Garcia, M.S., Morrell, J.I., and Pfaff, D.W. (1987). Distribution of luteinizing hormone-releasing hormone in the nervus terminalis and brain of the mouse detected by immunocytochemistry. J Comp Neurol. 255(2), 231–244.

Shang, J., Wan, Y., Luo, C., Ye, G., Geng, Q., Auerbach, A., and Li, F. (2020). Cell entry mechanisms of SARS-CoV-2. Proc Natl Acad Sci USA, 117(21), 11727–11734.

Solomon, T. (2021). Neurological infection with SARS-CoV-2 - the story so far. Nat Rev Neurol. 1–2.

Thakur, K.T., Miller, E.H., Glendinning, M.D., Al-Dalahmah, O., Banu, M.A., Boehme, A.K., et al. (2021). COVID-19 neuropathology at Columbia University Irving Medical Center/New York Presbyterian Hospital. Brain, awab148. Advance online publication. https://doi.org/10.1093/brain/awab148.

Tikellis, C., and Thomas, M.C. (2012). Angiotensin-Converting Enzyme 2 (ACE2) Is a Key Modulator of the Renin Angiotensin System in Health and Disease. Int J Pept. 2012:256294.

Ubuka, T., Moriya, S., Soga, T., and Parhar, I. (2018). Identification of Transmembrane Protease Serine 2 and Forkhead Box A1 As the Potential Bisphenol A Responsive Genes in the Neonatal Male Rat Brain. Front Endocrinol, 9, 139.

Ueha, R., Kondo, K., Kagoya, R., Shichino, S., Shichino, S., and Yamasoba, T. (2021). ACE2, TMPRSS2, and Furin expression in the nose and olfactory bulb in mice and humans. Rhinology, 59(1), 105–109.

van Riel, D., Verdijk, R., and Kuiken, T. (2015). The olfactory nerve: a shortcut for influenza and other viral diseases into the central nervous system. J Pathol. 235(2), 277–287.

von Bartheld, C.S. (2004). The terminal nerve and its relation with extrabulbar "olfactory" projections: lessons from lampreys and lungfishes. Microsc Res Tech. 65(1-2), 13–24.

Winkler, E. S., Bailey, A. L., Kafai, N. M., Nair, S., McCune, B. T., Yu, J., et al. (2020). SARS-CoV-2 infection of human ACE2-transgenic mice causes severe lung inflammation and impaired function. Nat Immunol. 21(11), 1327–1335.

Wirsig, C.R., and Leonard, C.M. (1986). Acetylcholinesterase and luteinizing hormone-releasing hormone distinguish separate populations of terminal nerve neurons. Neuroscience, 19(3), 719–740.

Ye, Q., Zhou, J., Yang, G., Li, R.-T., He, Q., Zhang, Y., et al. (2020). SARS-CoV-2 infection causes transient olfactory dysfunction in mice. bioRxiv [Preprint] doi: https://doi.org/10.1101/2020.11.10.376673. (accessed on May 30, 2021).

Zhang, A.J., Lee, A.C., Chu, H., Chan, J.F., Fan, Z., Li, C., et al. (2020). SARS-CoV-2 infects and damages the mature and immature olfactory sensory neurons of hamsters. Clin Infect Dis. 15:ciaa995. doi: 10.1093/cid/ciaa995

Zhao, M.M., Yang, W.L., Yang, F.Y., Zhang, L., Huang, W.J., Hou, W., et al. (2021). Cathepsin L plays a key role in SARS-CoV-2 infection in humans and humanized mice and is a promising target for new drug development. Signal Transduct Target Ther. Mar 27;6(1):134.

Zheng, J., Wong, L.R., Li, K., Verma, A.K., Ortiz, M., Wohlford-Lenane, C., et al. (2021). COVID-19 treatments and pathogenesis including anosmia in K18-hACE2 mice. Nature 589, 603–607.

Zhou, B., Thao, T.T.N., Hoffmann, D., Taddeo, A., Ebert, N., Labroussaa, F., et al. (2020). SARS-CoV-2 spike D614G variant confers enhanced replication and transmissibility. bioRxiv [Preprint]. doi: 10.1101/2020.10.27.357558. (accessed on May 30, 2021).

Zubair, A.S., McAlpine, L.S., Gardin, T., Farhadian, S., Kuruvilla, D.E., and Spudich, S. (2020). Neuropathogenesis and Neurologic Manifestations of the Coronaviruses in the Age of Coronavirus Disease 2019: A Review. JAMA Neurol. 77(8), 1018–1027.

